# Brain structures associated with reading and their abnormalities in dyslexia: a whole-brain analysis

**DOI:** 10.1101/2020.03.27.011577

**Authors:** T. Kujala, A. Thiede, P. Palo-oja, P. Virtala, M. Laasonen, J. Numminen, A. J. Sihvonen

## Abstract

Developmental dyslexia (DD) is a highly prevalent neurodevelopmental disorder, which often has a devastating influence on the individual’s academic achievement and career. Research on the neural origins of DD has continued for half a century, yielding, however, inconsistent results. The current study was set out to determine abnormalities of grey and white matter volumes in adults with DD and to shed light on neural architectures associated with reading and related skills. To this end, we conducted a whole-brain voxel based morphometry following current recommendations on analysis approaches, coupled with rigorous neuropsychological testing, to characterize the associations between neuroanatomy and skills vital for reading in DD. We found decreased volumes of grey matter in DD, comprising a left-hemispheric network including superior temporal and inferior frontal gyri, insula, the limbic system, and basal ganglia, and white matter, including the right middle temporal gyrus and hippocampus, as well as the right precuneus. These results are both consistent with the most robust previous findings on cortical abnormalities in DD and yield novel insight to the role of subcortical structures in DD, scarcely studied so far. Crucially, areas with decreased grey matter in DD overlapped with brain areas associated with technical reading skills. This supports the conclusion that the grey matter regions that we identified to have a low volume in DD are associated with the core areas vital for reading.

## 1. Introduction

Developmental dyslexia (DD) is a reading-skill impairment, which may emerge irrespective of adequate intelligence and reading instruction (Manual of Mental Disorders, 2000). The prevalence of dyslexia ranges between 5-17,5% (Shaywitz, 1998), which makes it the most common neurodevelopmental disorder. Due to its high prevalence and devastating influences on the individual’s academic achievements, career, self-esteem, and coping in the modern society, it is pertinent to understand the neural basis of DD. Yet, this task is very challenging due to the heterogeneity of its geno- and phenotype (Kere et al., 2014; McArthur et al., 2013; Zoubrinetzky et al., 2014) and the complexity of the neural network underlying reading (Kujala et al., 2007).

According to functional imaging studies, the key neural network for reading comprises frontotemporoparietal circuits predominantly in the left hemisphere (e.g., Levy et al., 2009; Welcome and Joanisse, 2012). During word reading, the early visual processing occurs in the left inferior occipitotemporal (OT) cortex (Stoodley and Stein, 2013). The OT along with cerebellum are the starting points for the two major forward-driving nodes in the network of reading (Kujala et al., 2007). After the left OT areas, reading involves left parietal cortex and left inferior frontal gyrus (IFG; Levy et al., 2009).

The endeavor to find anomalies in the reading circuitry in DD to reveal its neural basis has continued for over 50 years (e.g., Drake, 1968; Ramus et al., 2018). However, the previous morphological studies, including those utilizing modern neuroimaging methods, have offered relatively few replicated results on the neural basis of DD and the association between neuroanatomy and skills pertinent for reading.

The current study employed a rigorous neuropsychological testing of cognitive functions vital for reading and voxel-based morphometry (VBM), an automated MRI method for assessing focal brain changes (Ashburner and Friston, 2000). Since its introduction, VBM has become a standard method to analyze neuroanatomical abnormalities in various disorders, including DD with over 20 published studies (Brambati et al., 2004, Brown et al., 2001, Dole et al., 2013, Eckert et al., 2016, Eckert et al., 2005, Evans et al., 2014, Hoeft et al., 2007, Jednoróg et al., 2015, Jednoróg et al., 2014, Krafnick et al., 2014, Kronbichler et al., 2008, Menghini et al., 2008, Pernet et al., 2009a, Pernet et al., 2009b, Silani et al., 2005, Siok et al., 2008, Steinbrink et al., 2008, Tamboer et al., 2015, Vinckenbosch et al., 2005, Xia et al., 2016, Yang et al., 2016). Whereas the results have been spatially discorded in DD, the most frequently reported brain regions in VBM findings include left posterior temporal and temporoparietal areas, but with both increased and decreased grey matter (GM) volume reported in DD (e.g., Brambati et al., 2004; Silani et al., 2005; Hoeft et al., 2007). In addition, DD has been associated with reduced GM (both modulated and non-modulated data) bilaterally in the frontal lobe (Brown et al., 2001) as well as in the left superior frontal gyrus and IFG (Brown et al., 2001; Brambati et al., 2004), bilateral OT regions (Brambati et al., 2004; Eckert et al., 2005; Kronbichler et al., 2008), subcortical structures (caudate and thalamus, Brown et al., 2001), and cerebellum (Brown et al., 2001; Brambati et al., 2004; Eckert et al., 2005; Kronbichler et al., 2008). However, a number of studies with no significant morphological findings in DD have also been published (e.g., Pernet et al., 2009a, b; Casper et al., 2018).

Reduced WM volumes in DD have been reported in left-hemispheric frontal areas, post-central gyrus, paracentral lobule, and temporo-parietal region (Eckert et al., 2005; Silani et al., 2005). Furthermore, WM volume reductions have been reported subcortically, in striatum and hippocampus (Wang et al., 2019), and with a matched-brain morphometry approach, in corona radiata, internal capsule (Eckert et al., 2017).

Three recent meta-analyses have tackled with the heterogeneous morphometric findings in DD. In 2013, Richlan and colleagues evaluated nine VBM studies on DD and found reduced GM in the superior temporal areas bilaterally in DD (Richlan et al., 2013). A few years later, Eckert and colleagues evaluated 11 VBM studies on DD, and reported lower GM volume (i.e. modulated data; adjusted for total GM volume) in left superior temporal sulcus, left orbitofrontal cortex, and the right cerebellum in participants with DD (Eckert et al., 2016). Interestingly, one of the most robust findings is reduced total brain volume in subjects with DD compared with typical readers, a finding confirmed by a meta-analysis including 1164 participants across 18 studies (Ramus et al., 2018). However, the direction of association between the reduced brain volume and DD, let alone its possible aetiological factor for DD remain currently unknown.

Overall, there is enormous variation between the results obtained on the structural brain anomalies in DD, which presumably partly results from the heterogeneity of DD, but also has raised the concern of their reliability (Ramus et al., 2018). Across the published studies, a mixed selection of statistical thresholding and corrections for multiple tests have been applied, if any. When voxel-based neuroimaging methods (e.g. VBM) are used to reliably identify the neuroanatomical changes in DD, a combination of reasonable cluster-size threshold and voxel extent is needed to produce a desirable balance between Types I and II error rates (Lieberman and Cunningham, 2009). In addition, most of the studies consist of small sample sizes, with only few studies including more than 16 dyslexics (see Eckert et al., 2016, for a meta-analysis). Furthermore, consistency in adjusting the morphological analyses for confounding effects is lacking. Another core problem is the bias for publishing studies reporting group differences. Due to these reasons, diving the more robust findings from the less reliable is challenging (see Ramus et al., 2018, for a review).

To attain reliable VBM results, it is critical how the preprocessing of MRI data is carried out, in other words, how the GM and WM probabilistic maps are formed. As in any voxel-based imaging analysis, following published guidelines, ensuring efficient registration, as well as controlling for factors that affect brain size (e.g. age) aid to achieve dependable results and extra sensitivity (Barnes et al., 2010; Ashburner and Friston, 2001; Pell et al., 2008; Li X et al., 2013; Da Ma et al., 2018). One crucial preprocessing step is modulation, which allows testing for regional differences in the absolute amounts (volume) of GM or WM (Ashburner and Friston, 2000; Good et al., 2001). Ramus et al. (2018) published a well-founded list of methodological recommendations for more reliable research on the neuroanatomy of DD and encouraged to systematically use relevant covariates to diminish their confounding effects on between-subject comparisons. In general, it is strongly recommended to include an adjustment for head size and other nuisance variables like gender and age in the morphological analyses (Barnes et al., 2010; Da Ma et al., 2018; Li X et al., 2013; Pell et al., 2008). Moreover, other seldom controlled variables in DD analyses are verbal and full-scale intelligence quotients (IQ), which were suggested to be correlated with brain size (McDaniel, 2005).

Based on this information, and following the previously published recommendations (Ramus et al., 2018), we set out to evaluate the GM and WM volume abnormalities in adult dyslexic individuals, whose reading-skill and neuropsychological profiles were rigorously assessed with a carefully composed test battery. We investigated brain properties while systematically using relevant covariates in the analyses. Furthermore, the connection between regional GM and WM volume and reading-related skills were analyzed as well as the association between these skills and group findings.

## 2. Materials and methods

### 2.1 Procedure

The data were collected in three separate sessions. In the first session neuropsychological testing was carried out. The second session included functional imaging (combined magneto- and electroencephalography, the data from which will be reported elsewhere) and responding to questionnaires. In the third session, anatomical MRI data were acquired.

### 2.2 Participants

Participants were recruited via social media, website of a related project, and through a local learning disabilities association. Forty-five right-handed Finnish-speaking participants completed the MR-imaging, the final sample consisting of 22 typically reading and 23 dyslexic participants. The groups were balanced in age, years of education, years of music education (since it may improve language skills; see, e.g., Kraus and Chandrasekaran, 2010), and sex (see Table 1 for descriptive statistics). The groups showed no significant differences in the demographic variables, whereas they significantly differed in the composite scores of phonological processing, technical reading, and working memory (Table 1). However, they differed in the full-scale IQ (FIQ), which was, therefore, used as a covariate in the analyses.

**Table 1.** Neuropsychological tests and composites (bold).

| Neuropsychological composites (bold)<br>and individual tests | Median (IQR) | | $p_{corr}$ | signif.<br>level |
| --- | --- | --- | --- | --- |
| | Dyslexic ($n = 23$) | Control ($n = 22$) | | |
| **Phonological processing [z]** | -0.2 ( 1.2 ) | 0.4 ( 0.4 ) | <.000 | *** |
| Pig Latin (accuracy) | 9.0 ( 7.0 ) | 15.0 ( 1.0 ) | <.000 | *** |
| Nonword span length (accuracy) | 12.0 ( 3.0 ) | 13.0 ( 4.0 ) | .083 | ns |
| Rapid Alternate Stimulus naming<br>(RAS) (speed of second trial) | 30.0 ( 10.7 ) | 24.0 ( 6.4 ) | <.000 | *** |
| **Reading, technical [z]** | -0.3 ( 0.9 ) | 0.6 ( 0.2 ) | <.000 | *** |
| word list reading (accuracy) | 30.0 ( 1.0 ) | 30.0 ( 0.0 ) | .005 | ** |
| word list reading (speed) | 31.0 ( 11.4 ) | 19.3 ( 2.9 ) | <.000 | *** |
| pseudoword list reading (accuracy) | 21.0 ( 8.5 ) | 28.5 ( 3.5 ) | <.000 | *** |
| pseudoword list reading (speed) | 72.9 ( 32.6 ) | 40.1 ( 7.7 ) | <.000 | *** |
| text reading (accuracy) # | 98.2 ( 1.1 ) | 99.4 ( 0.8 ) | <.000 | *** |
| text reading (speed) # | 305.0 ( 67.0 ) | 449.0 ( 62.8 ) | <.000 | *** |
| **Full Intelligence Quotient** | 104.5 ( 17.3 ) | 118.0 ( 11.7 ) | <.000 | *** |
| Verbal IQ [Wechsler Adult Intelligent<br>Scale (WAIS)-III Similarities and<br>Vocabulary] | 103.0 ( 20.0 ) | 115.0 ( 10.0 ) | <.000 | *** |
| Performance IQ (WAIS-III Block<br>design and Matrix reasoning) | 113.0 ( 11.0 ) | 120.5 ( 11.6 ) | .004 | ** |
| **Working memory functions** | 19.0 ( 7.5 ) | 24.0 ( 5.8 ) | .007 | ** |
| WMS-III Number series | 10.0 ( 3.5 ) | 13.0 ( 3.8 ) | <.000 | *** |
| WMS-III Visual Series | 9.0 ( 5.0 ) | 10.5 ( 3.0 ) | .193 | ns |

Notes. Group sizes (n) and median values of all variables in the Dyslexic and Control groups with interquartile range (IQR) in parentheses. Group differences were tested with Wilcoxon sign-rank test, and significance levels of FDR-corrected $p$ -values are indicated by asterisks. ns – non-significant. #– Not included in the technical reading composite score.

The participant was classified as dyslexic if 1) a recent statement on dyslexia diagnosis was available from a health-care professional (e.g., psychologist), or the participant had both 2) reading-related problems in childhood as evaluated with the adult ARHQ and confirmed in an interview, and 3) a performance of at least one standard deviation (SD) below the average of age-matched standardized control data (Laasonen et al., 2010) in at least two different reading subtests (word list reading, pseudoword list reading, text reading) in either speed or accuracy (Table 1). Participants classified to the Control group 1) had no language-related problems (and neither did their parents nor siblings), 2) reported no childhood problems in reading or writing as indicated by the ARHQ, and 3) performed within norm in at least two reading subtests in both speed and accuracy.

The exclusion criteria were as follows (self-reported except for IQ, which was tested): attention deficit evaluated by the Adult ADHD Self-Report Scale ASRS-v1.1 questionnaire (Kessler et al., 2005), developmental language disorder or other language impairments, other neurological or psychiatric disorders, substance abuse, medication affecting central nervous system, uncorrected hearing or visual deficit, left-handedness, an individualized school curriculum, early bilingualism, and a performance IQ below 80. MRI data of four participants could not be obtained due to non-detachable metal in the body or pregnancy (three dyslexics, one control).

The experiments were undertaken with the understanding and written consent of each subject, and following the Code of Ethics of the World Medical Association (Declaration of Helsinki). The Coordinating Ethics Committee of The Hospital District of Helsinki and Uusimaa approved the study protocol.

### 2.3 Neuropsychological tests and questionnaires

Participants filled out questionnaires concerning background information, DD in family, Adult Reading History Questionnaire (ARHQ; Lefly & Pennington, 2000), and Attention-Deficit/Hyperactivity Disorder (ADHD) Self-Report Scale Symptom Checklist ASRS-v1.1 (Kessler et al., 2005). The questions on the background included the participants’ language skills and usage; language difficulties, neurological, psychiatric, and hearing disorders of him/her and the relatives (parents and siblings); the participant’s other health issues and vision; music education and listening; and education and employment status. One questionnaire assessed the family history of DD.

The neuropsychological test battery was designed to assess IQ, reading, phonological processing (phonological awareness, phonological short-term memory and rapid access of phonological information; Torgesen et al., 1994), and working memory functions (Table 2). The verbal IQ was evaluated with the subtests Similarities and Vocabulary, and the performance IQ was assessed with the subtests Block Design and Matrix Reasoning from the Wechsler Adult Intelligence Scale III (Wechsler, 2005). Technical reading skills (accuracy and speed, Cronbach’s α = .87) were tested with word and pseudoword list reading (Nevala et al., 2006). The domain of phonological processing (Cronbach’s α = .69) included neuropsychological tests as follows: ‘Pig Latin’ test for assessing phonological awareness (Nevala et al., 2006; the participant is asked to change the first syllables between two heard pseudowords and produce aloud the new pseudowords, e.g., kouta-mesi -> meuta kosi), non-word span length for phonological short-term memory (Laasonen et al., 2002; the participant repeats lengthening sequences of heard pseudowords, e.g., pola-sine-heka), and rapid alternating stimulus naming for rapid serial naming (Wolf, 1986; the participant names as rapidly and accurately as possible a 10×5 matrix of alternating colors, letters, and numbers). Working memory functions were evaluated with Letter-number Series and Visual series subtests from the Wechsler Memory Scale, WMS-III (Wechsler, 2008).

**Table 2.** Demographic and morphological data *Group sizes (n) and mean values of background variables in the Dyslexic and Control groups with standard deviation in parentheses. P-values show Chi Squared (χ^2^), and independent-samples t-test (t) statistics for group comparisons*.

|  | **Dyslexic**<br>**(n = 23)** | **Control**<br>**(n = 22)** | **P value** |
| --- | --- | --- | --- |
| **Demographic** |  |  |  |
| Gender (male/female) | 11/12 | 10/12 | 1.000<br>($\chi^2$) |
| Age (years) | 31.3 (8.6) | 29.8 (5.9) | .530 (t) |
| Education (years) | 15.7 (5.2) | 16.1 (4.4) | .817 (t) |
| Musical education (years) | 3.0 (7.8) | 3.7 (5.5) | .730 (t) |
| **Morphological** |  |  |  |
| Grey matter volume<br>(litres) | 0.8 (0.1) | 0.8 (0.1) | .705 (t) |
| White matter volume<br>(litres) | 0.5 (0.1) | 0.5 (0.1) | .680 (t) |
| Cerebrospinal fluid<br>volume (litres) | 0.5 (0.1) | 0.5 (0.2) | .817 (t) |
| Total intracranial volume<br>(litres) | 1.8 (0.3) | 1.8 (0.3) | .934 (t) |
| Total brain volume<br>(litres) | 1.3 (0.1) | 1.3 (0.1) | .666 (t) |

We chose to use composite scores instead of the individual variables of single tasks for two reasons: in order to reduce the number of analyses and to reduce the error variance related to single task performance. Unfortunately, the size of the data in the current study did not allow for conducting a factor analysis over the variables and, thus, were chose to use the classifications based on previous theoretical and factor-analytic studies but also checked the internal consistency of our domain variables with Cronbach’s α (see above). Composite scores of the test results (bolded in Table 2) were formed for phonological processing and technical reading by converting the raw scores (of subtests listed in Table 2 below the respective composite) to *z*-scores and averaging them, and for working memory the composite was formed according to WMS-III (Wechsler, 2008).

### 2.4 MRI data acquisition

Participants were scanned with a 3T MRI Siemens Skyra scanner (Siemens Healthcare, Erlangen, Germany) using a 32-channel head coil, at AMI center in Aalto University, Finland. High-resolution magnetization prepared rapid acquisition gradient-recalled (MPRAGE) T1 images were obtained (flip angle = 7°, TR = 2530 ms, TE = 3.3 ms, voxel size = 1.0 × 1.0 × 1.0 mm^3^).

### 2.5 Data analysis

#### 2.5.1 Voxel-based morphometry

Morphometric analysis was carried out using VBM (Ashburner and Friston, 2000) and the Statistical Parametric Mapping software (SPM8, Wellcome Department of Cognitive Neurology, UCL) under MATLAB 8.0.0 (The MathWorks Inc., Natick, MA, USA, version R2014b). VBM is an MRI analysis technique that allows comparison of GM and WM differences in focal brain regions between groups (Ashburner and Friston, 2000). After reorienting the individual T1 images using the anterior commissure as a landmark for the origin, Unified Segmentation (Ashburner and Friston, 2005) with medium regularization was applied to the T1 images, segmenting them precisely into GM, WM, and cerebrospinal fluid probability maps before normalizing them into the Montreal Neurological Institute (MNI) space using SPM8 normalization. To preserve the original signal strength during the normalization, GM and WM probability maps were modulated. After this, to reduce residual inter-individual variability, GM and WM probability maps were smoothed using an isotropic spatial filter (FWHM = 6 mm). During each step, the images were visually checked for potential registration errors. Modulated probability maps were used to calculate TIV (volumes of GM, white matter volume and cerebrospinal fluid added together), TBV (GM volume and white matter volumes added together) (Malone et al., 2015).

#### 2.5.2 Statistical analyses

Preprocessed GM and WM images were then entered into a second-level analysis. First, three one-sample t-tests including all subjects (N=45) were calculated to evaluate the focal GM and WM structures associated with better performance in technical reading, phonological, and working memory tests. Then, using independent-sample t-tests, two different contrasts (Controls>Dyslexics, Dyslexics>Controls) were calculated to evaluate the GM and WM volumetric differences between the dyslexics and controls. All results were corrected for nonstationarity (Hayasaka et al., 2004) and spmT-maps were thresholded at a whole-brain uncorrected P<.005 threshold and a familywise error rate (FWE) corrected P<.05 at the cluster level, a combination which has been shown to produce a desirable balance between Types I and II error rates, comparable to false discovery rate (FDR)(Lieberman and Cunningham, 2009), and corrected for non-isotropic smoothness using VBM8 toolbox (http://dbm.neuro.uni-jena.de/vbm8/VBM8-Manual.pdf). Exact neuroanatomical regions were identified using the Automated Anatomical Labeling Atlas (Tzourio-Mazoyer et al., 2002) included in the xjView toolbox (http://www.alivelearn.net/xjview/).

Partial correlations (two-tailed) were calculated between each individual significant cluster and the three composite scores (technical reading, phonological processing, working memory; see Table 1) using SPSS (IBM Corp. Released 2012. IBM SPSS Statistics for Windows, Version 24.0. Armonk, NY: IBM Corp.). To control for multiple comparisons, FDR approach was used and only significant results are reported.

All statistical analyses were adjusted for age, gender, and TIV (Barnes et al., 2010; Da Ma et al., 2018; Li X et al., 2013; Pell et al., 2008). Furthermore, to follow the previously published recommendations (Ramus et al., 2018) and to take into account the group difference, FIQ was also added as a covariate.

## 3. Results

### 3.1 Anatomical correlates of neuropsychological tests

We first determined how reading, phonological processing, and working memory were associated with brain structures including both groups in the analysis (Fig. 1; Table 3). Better performance in technical reading (accuracy and speed combined) was associated with greater GM volume in a cluster comprising the left temporal (STG and middle temporal gyrus (MTG), fusiform gyrus), limbic (amygdala, hippocampus, parahippocampal gyrus) and insular areas (R=.634, P<.001). In addition, better technical reading performance was associated with greater WM volume in one cluster comprising right frontal areas (IFG and precentral gyrus), basal ganglia (putamen, globus pallidus), insula, pons, parahippocampal gyrus, and left cerebellum (R=.606, P<.001). Higher working memory scores were associated with greater WM volume in the brainstem and bilaterally in the cerebellum (R=.518, P=.001).

**Figure 1.**
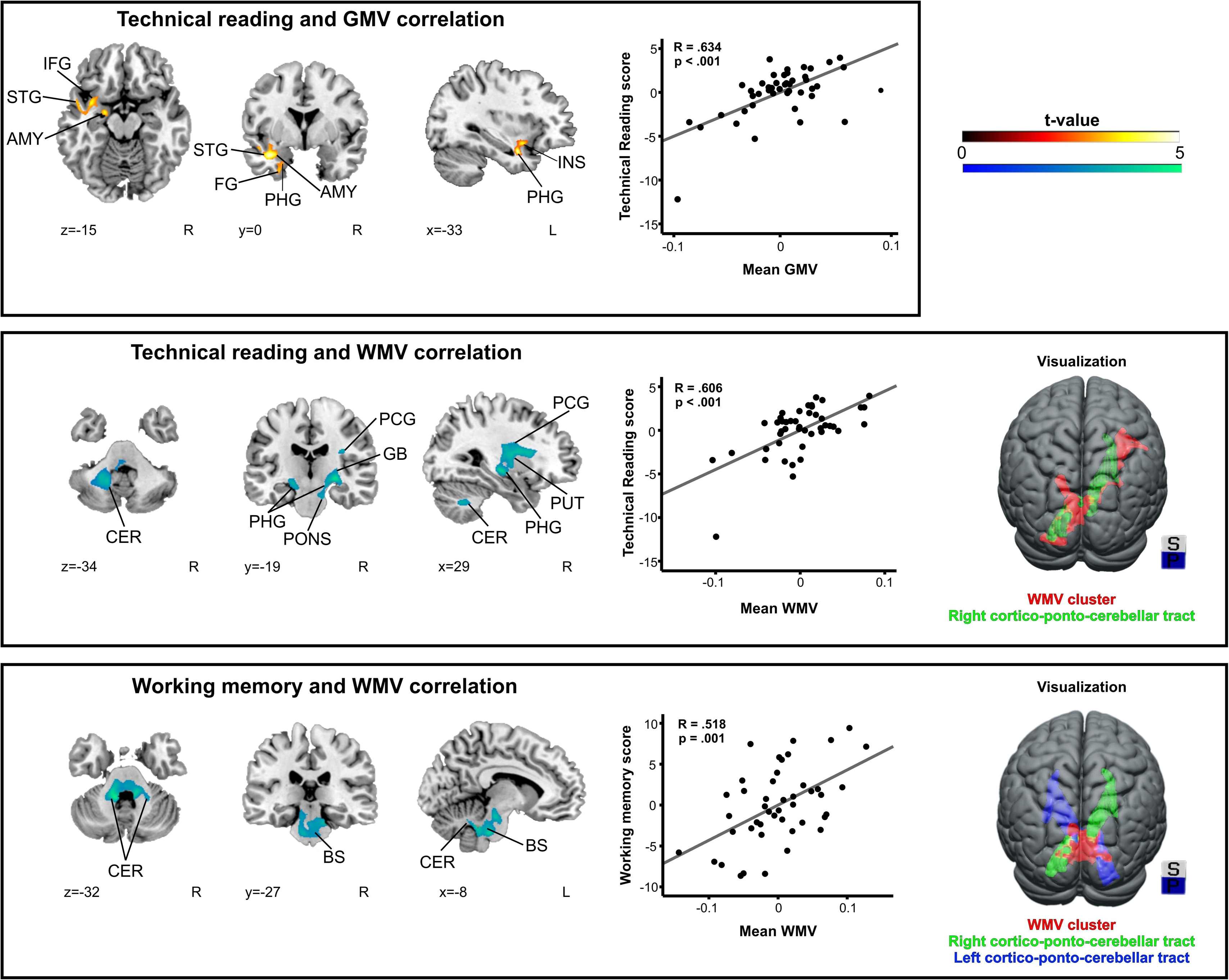
Grey and white matter volume associations with reading-related skills (see also Table 3). Top: Grey matter volume and technical reading score; Middle: White matter volume and technical reading score; Bottom: White matter volume and working memory score. N = 45. Neurological convention is used with MNI coordinates at the bottom left of each slice. All statistical maps are thresholded at a cluster-level FWE-corrected p < 0.05 threshold and corrected for nonstationarity. Grey and white matter volume correlations to reading-related skills are shown with scatter plots. For visualization purposes, a white matter atlas (https://www.natbrainlab.co.uk/atlas-maps) was used to present the white matter volume clusters. AMY = amygdala, BS = brainstem, CER = cerebellum, FG = fusiform gyrus, GB = globus pallidus, GMV = grey matter volume, IFG = inferior frontal gyrus, L = left, P = posterior, PCG = precentral gyrus, PHG = parahippocampal gyrus, R = right, S = superior, STG = superior temporal gyrus, WMV = white matter volume.

**Table 3.**
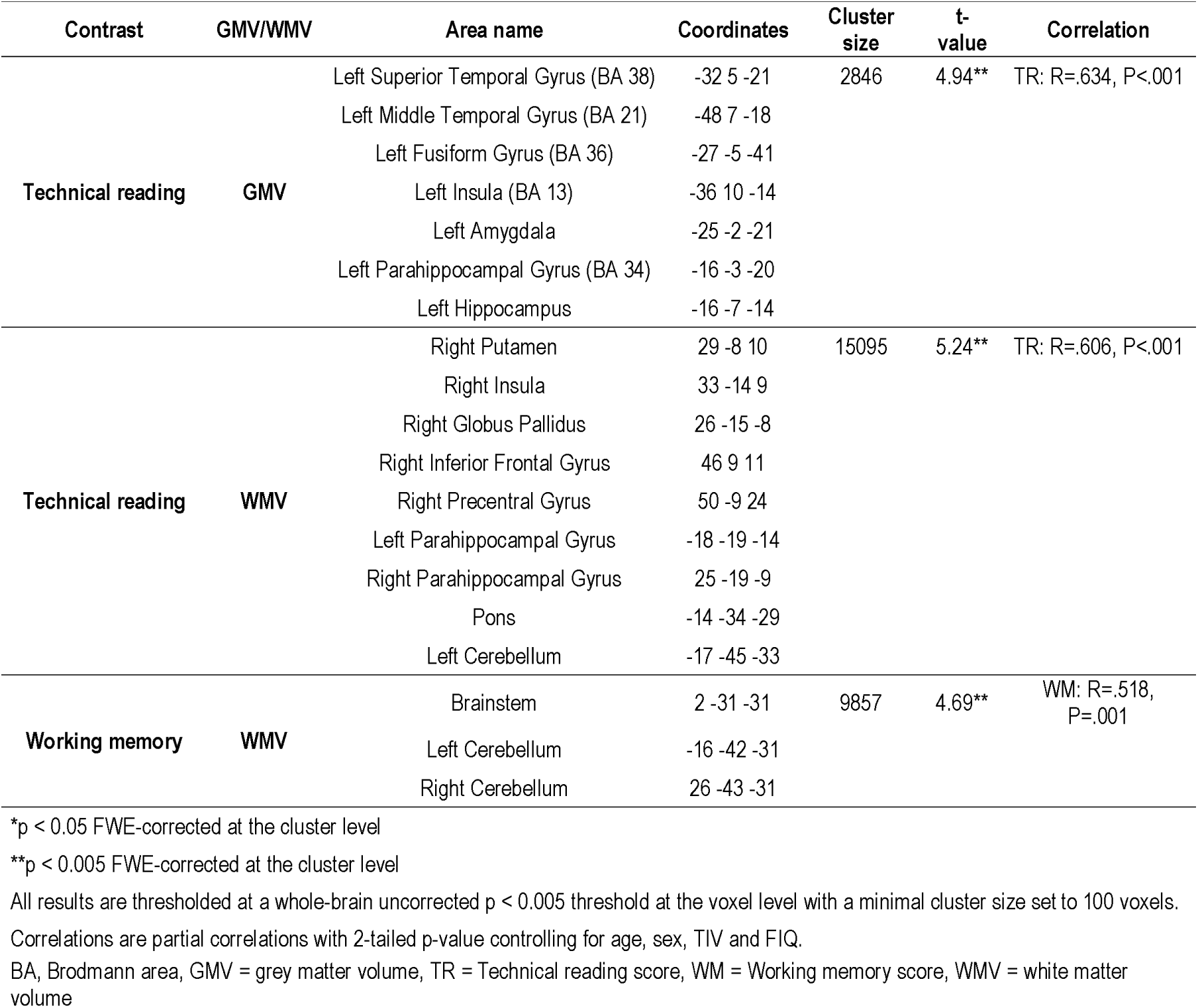
Focal grey and white matter volume associations with the neuropsychological composite scores.

When the two significant WM clusters (i.e. technical reading and working memory) were overlaid with white matter atlas (https://www.natbrainlab.co.uk/atlas-maps) for visualization purposes, both bore a resemblance to cortico-ponto-cerebellar (CPC) tracts (Fig. 1). We carried out an exploratory *post-hoc* analysis evaluating the amount of overlap between the two WM clusters and the CPC tracts. From technical reading WM cluster, 24.4% situated in the right CPC and from the working memory WM cluster, 23.7% and 33.5% situated in the left and right CPC pathway, respectively.

### 3.2 Neuroanatomical group differences

First, we evaluated whether the controls and dyslexics had significant brain volumetric differences in TIV, total brain volume, GM, WM or cerebrospinal fluid. Five independent-sample *t*-tests were performed, out of which none were statistically significant. The groups showed no significant differences in TIV t(43)=.083, P=.934, total brain volume t(43)=.434, P=.666, GM volume t(43)=.381, P=.705, WM volume t(43)=.415, P=.680 and cerebrospinal fluid t(43)=-.232, P=.817.

In whole-brain analyses comparing the groups, greater GM volume in Controls than Dyslexics was found in a cluster comprising the left superior temporal gyrus (STG), IFG, insula, as well as in the left limbic (amygdala, hippocampus, subcallosal gyrus) and basal ganglia areas (claustrum, putamen; Fig. 2 and Table 4). The mean GM volume in the observed areas correlated significantly with technical reading score (R=.432, P=.015). Crucially, there was a significant overlap between the aforementioned group difference and the GM structures which correlated with technical reading skills (Fig. 2 and Table 4). The overlapping area comprised the left STG, insula, and limbic areas (amygdala, hippocampus), and greater GM volume in these areas correlated with higher technical reading score (R=.575, P<.001).

**Figure 2.**
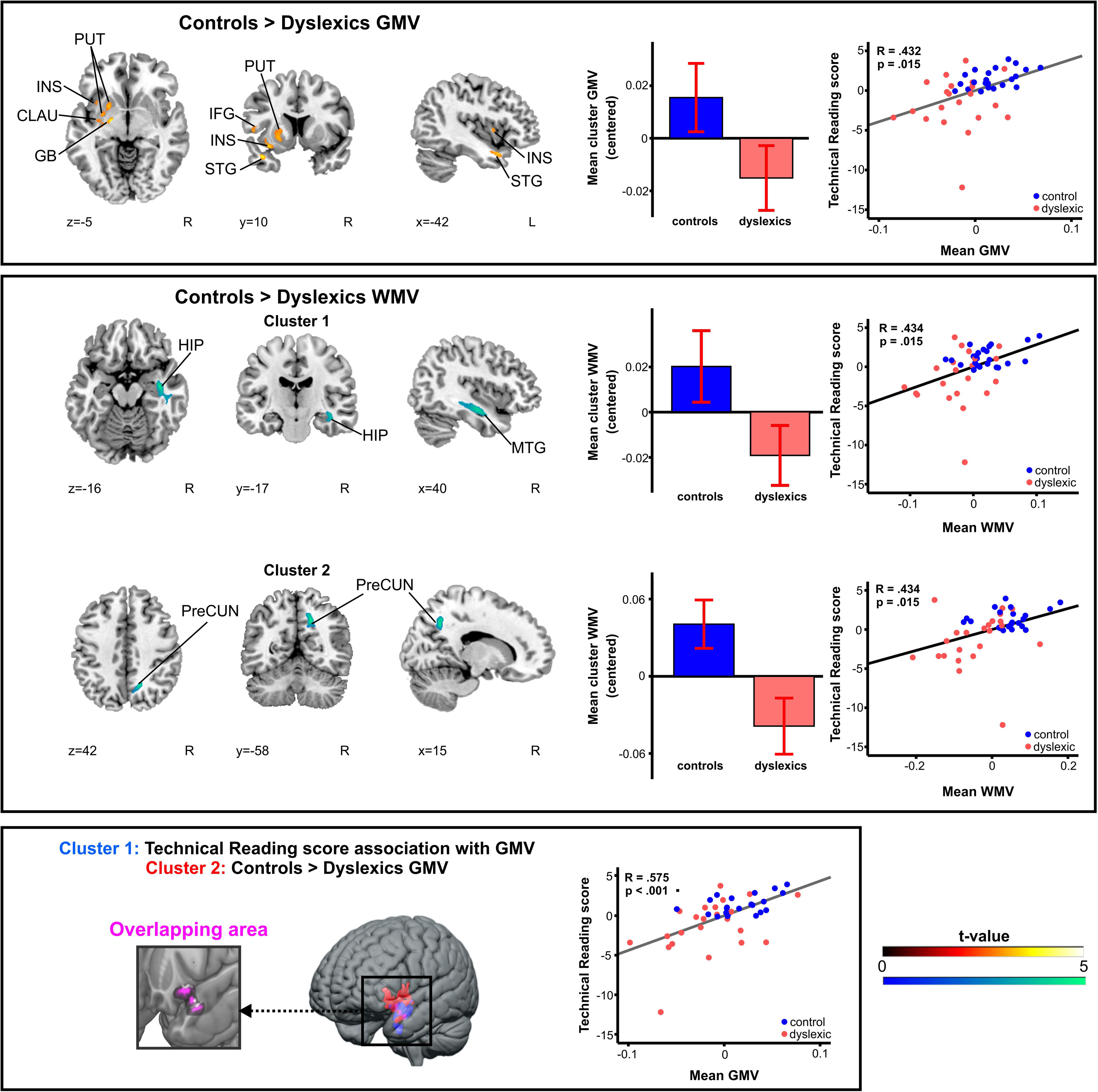
Grey and white matter volume group differences (see also Table 4). Top: Controls > Dyslexics, grey matter volume; Middle: Controls > Dyslexics, white matter volume; Bottom: Overlap image (purple) of Controls > Dyslexics, grey matter volume (red) and grey matter volume and technical reading score correlation analysis (blue). N = 45. Neurological convention is used with MNI coordinates at the bottom left of each slice. All statistical maps are thresholded at a cluster-level FWE-corrected p < 0.05 threshold and corrected for nonstationarity. Grey and white matter volume correlations to reading-related skills are shown with scatter plots. Bar plots for mean GMV in significant clusters (Table 4) are shown: bar = mean (centered), error-bar = standard error of mean. CLAU = claustrum, GB = globus pallidus, GMV = grey matter volume, HIP = hippocampus, IFG = inferior frontal gyrus, INS = insula, L = left, MTG = middle temporal gyrus, PUT = putamen, PreCUN = precuneus, R = right, STG = superior temporal gyrus, WMV = white matter volume.

**Table 4.**
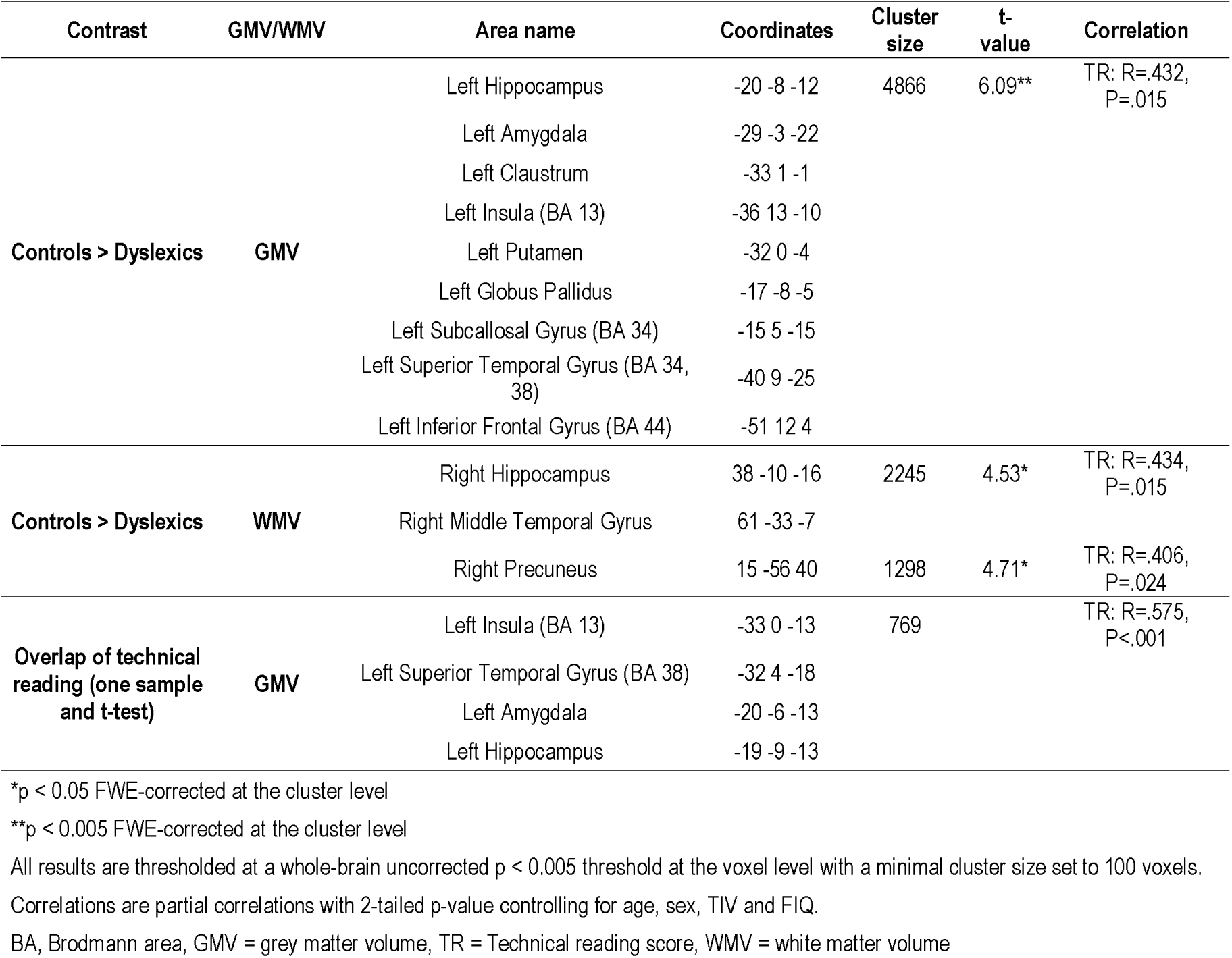
Grey and white matter volume differences between the dyslexic and control groups.

In addition, the control participants had greater WM volume in two clusters in the right MTG and hippocampus as well as in the right precuneus compared with the dyslexic participants (Fig. 2 and Table 4). Again, greater WM volume in these areas correlated significantly with a higher technical reading score (R=.434, P=.015 and R=.406, P=.024, respectively).

## 4. Discussion

There is an obvious need to understand neural underpinnings of DD, which is highly prevalent and can have devastating effects on the individual affected, yet, no clear consensus has so far been reached (e.g., Ramus et al., 2018, for a review). Our study was set out at illuminating the neuroanatomical basis of DD using a whole-brain volumetric analysis and a thorough reading and neuropsychological test battery. Our main finding was a decreased volume of GM in participants with DD, comprising left frontotemporal and limbic regions as well as the left basal ganglia, a cluster in which greater GM volume was associated with better technical reading abilities across the whole sample. The overlap of areas correlating with reading skills with those having a reduced volume in participants with DD provides a firm basis to interpret that these regions represent the neural origins of dyslexia. These include areas (left STG and IFG, insula) that have previously been linked with the reading network (Kujala et al., 2007) and functional and anatomical abnormalities in dyslexia (Richlan et al., 2009; 2013). Moreover, our study unravels subcortical structures, the involvement of which in reading and dyslexia have previously been scarcely studied. Also for these areas, our study found associations both with volumetric reductions in participants with DD and reading skills, completing the picture on the neural basis of dyslexia. Moreover, whereas lower total brain volume has so far been the most systematic finding in dyslexia (Ramus et al., 2018, for a review), the total brain volume of our dyslexic sample did not differ from that of our controls. This crucially suggests that dyslexia does not merely result from an overall lower brain volume but from changes in specific brain regions.

### 4.1 Associations between anatomy and reading-related skills

As could be expected, better performance in technical reading was significantly correlated with neural structures comprising both hemispheres, since a wide range of functions subserved by multiple brain areas are needed for fluent and accurate reading (e.g., Paulesu et al., 2000; Kujala et al., 2007; Levy et al., 2009; Welcome and Joanisse, 2012; Oberhuber et al., 2013). The GM volume findings included the left temporal (STG, MTG, fusiform gyrus), limbic (amygdala, hippocampus, parahippocampal gyrus), and insular areas. In addition, higher technical reading scores were associated with a larger WM volume in right frontal areas (IFG and precentral gyrus), basal ganglia (putamen, globus pallidus), insula, pons, as well as parahippocampal gyrus bilaterally and left cerebellum, closely resembling CPC pathways which have been implicated in language (for a review, see Vias and Dick, 2017). Overall, these results are compatible with a large body of previous research associating reading skills and subskills with brain structures in frontal, temporal, cerebellar, and subcortical areas (Paulesu et al., 2000; Kujala et al., 2007; Levy et al., 2009; Welcome and Joanisse, 2012; Oberhuber et al., 2013; Stoodley and Stein, 2013).

Higher working memory scores were associated with increased WM volume in the brainstem and bilaterally in the cerebellum. This is consistent with a previously found association between working memory load and the amplitude of brainstem responses (Sörqvist et al., 2012), indicating the involvement of brainstem in working memory functions. Brainstem also regulates vigilance (Iovino et al., 2019). Therefore, it could be speculated that these associations are related to attention, which is tightly involved in working memory functions requiring “attentional spotlight” to items that are actively consciously processed (e.g., Rhodes and Cowan, 2018). The association between the cerebellar WM volume and working memory is in accordance with currently known cerebellar functions. Cerebellum is involved in regulating the direction of attention, detecting errors, timing and sequencing, as well as in associative learning, all vitally involving working memory (Stoodley and Stein, 2013, for a review). This is also compatible with the rich cerebellar interconnections with the prefrontal cortex and other association cortices (Stoodley and Stein, 2013), which belong to the neural network responsible for working memory functions (Knight et al., 1999).

We found no significant correlations between phonological scores and neuroanatomical findings. A potential reason for this might be that only relatively small brain areas, suggested to be restricted to the left temporal and angular gyri (Glezer et al., 2016) process phonology and that the present analyses controlling for various nuisance variables impede small effects to become significant. Moreover, while both groups had more than 20 subjects, our overall sample size remained moderate, which in turn decreases power to observe small effects in the data. Furthermore, using our phonological composite score in the correlation analysis might not be optimal, since the tests included tap also into other functions besides pure phonological processing. Another possible explanation is that phonological deficits might primarily arise from subtle anatomical and functional changes in pathways interconnecting posterior and anterior superior temporal regions (Richardson et al., 2011) rendering the overall methodology (i.e. VBM) disadvantageous here.

### 4.2 Neuroanatomical group differences

We found reduced GM volume in dyslexic participants in left-hemispheric regions of STG, IFG, insula, limbic system, and basal ganglia, and WM volume in the right temporal, hippocampal and parietal areas. Importantly, the greater GM and WM volumes correlated with better technical reading skills, that is, reading accuracy and speed, directly linking the same brain areas to the core reading skills and reduced neural volumes in DD, which underlines the robustness and reliability of our results. However, interestingly, even though the most robust finding in previous studies has been a reduced total brain volume in DD (see Ramus et al., 2018, for a meta-analysis), we found no group differences in the brain, GM, or WM volumes. Thus, overall volumetric differences do not explain the deficits in reading and related skills in the current sample of dyslexics. This lack of group differences in gross brain volumetric measures is particularly interesting since it has been speculated that reduced brain volume in DD could either be associated with the etiology of DD or a consequence of this disorder (Ramus et al., 2018). However, contradicting results have been published (Frye et al., 2010) and the finding could depend on the population studied (adults vs. children). Our data showing no group differences in a range of volumetric brain measures (TIV, total brain volume, GM volume, WM volume or cerebrospinal fluid volume) suggest that the occurrence of DD may not (only) rely on brain volume reduction as a predisposing factor or as a de rigueur developmental consequence.

Our group and correlational GM volume results in STG, IFG, and insula are consistent with earlier neuroanatomical findings. For example, according to the meta-analysis of Richlan et al. (2013) the most consistent GM reductions in DD have been found in superior temporal areas and according to the meta-analysis of Ramus et al. (2018) in left perisylvian and OT regions. Furthermore, a large number of studies proposed that these areas belong to a network vital for reading (Kujala et al., 2007; Paulesu et al., 2000; Kujala et al., 2007; Levy et al., 2009; Welcome and Joanisse, 2012; Oberhuber et al., 2013). Our results are also compatible with functional imaging studies on DD, which have revealed hypoactivations in superior, middle, and inferior temporal areas as well as IFG in the left hemisphere (Richlan et al., 2009, for a meta-analysis). Next, we will inspect our findings on brain structures with lower volume in DD and association with technical reading skills in the light of the currently known functions of these areas.

**Superior temporal areas** were consistently found to be active during reading (Sandak et al., 2004; Richlan, 2014), and proposed to reflect semantic (Helenius et al., 1998; Halgren et al., 2002) and phonological analysis (Jobard et al., 2003). A restricted lesion to left STG was reported to result in pure word deafness, with the patient being able to read, write, and perceive nonspeech auditory stimuli but not to correctly process speech input (Maffei et al., 2017). The reduced GM in the STG of our dyslexic participants might, therefore, reflect a speech-specific deficit, consistent with one of the leading theories on DD according to which reading impairments in DD are largely based on poor speech sound representations (Peterson and Pennington, 2012; see, however, Ramus, 2014). This result is highly compatible with a meta-analysis based on which it was proposed that the phonological processing deficit is an endophenotype of DD (Snowling and Melby-Lervåg, 2016).

Activations of **left IFG** have been reported for words and legal pseudowords (pseudowords not violating the rules of the language) but not for consonant strings (Wilson et al., 2005; Cornelissen et al., 2009) or faces (Cornelissen et al., 2009). Its subregion BA 44 was found to be activated by both words and pseudowords, whereas subregion 45 was only activated by words (during a lexical decision task; Heim et al., 2005). These areas were also found to be involved in rapid learning of novel word-forms (Kimppa et al., 2015; 2018), an effect not found in dyslexic children (Kimppa et al., 2018). Furthermore, it was proposed that left IFG participates in nonlexical phonological grapheme-phoneme conversion during reading (Heim et al., 2005). Overall, these results, as well as a meta-analysis (Bookheimer 2002), suggest that left IFG regions are involved in word and phonological speech processing. The reduced left IFG GM in our dyslexic participants might, therefore, reflect their observed problems of word-form processing and learning (Kimppa et al., 2018), as well as deficits in phonological processing (e.g., Snowling and Melby-Lervåg, 2016) and grapheme-phoneme conversion (Snowling, 1980).

Whereas the involvement of cortical areas in reading and reading impairment has been extensively studied, less attention has been paid to **subcortical areas** so far (see Krishnan et al., 2016, for a review). Our study revealed a reduced GM volume in the dyslexic sample in limbic (hippocampus and amygdala) and basal ganglia (claustrum and putamen) areas. Similar findings were obtained in the left hemisphere of dyslexic males by Casanova et al. (2005), reporting anomalies in amygdala, hippocampus, putamen, and globus pallidus in their volumetric analysis. Furthermore, in dyslexic children, a successful reading intervention was found to result in increased GM volume in several brain areas including hippocampus (Krafnick et al., 2011). Hippocampus is known to have a central role in memory functions, particularly in memory consolidation (see Buzsáki and Moser, 2013, for a review). A reduced GM in the hippocampus of the dyslexic participants could be associated with their poor ability to “tune in” or form memory representations of novel repetitive stimuli (see Ahissar, 2007, for a review; see also Kimppa et al., 2018). This was suggested to influence the efficacy of short-term memory and underlie a wide range of difficulties associated with DD (Ahissar, 2007).

Amygdala, besides having a central role in processing fear-eliciting stimuli (Markowitsch, 1998) was also suggested to be involved in memory and reward networks and to process valence, salience, and stimulus relevance (Sander et al., 2003; Janak and Tye, 2015, for reviews), the left amygdala having a higher affinity to language than the right one (Markowitsch, 1998). Processing of relevance and salience, in turn, are strongly linked with attention functions, which are known to be dysfunctional in at least some subgroups of dyslexic individuals (Hari et al., 2001; see Krause, 2017, for a review). Despite our participants having been screened for ADHD with a questionnaire, it is possible that they had some attentional difficulties that do not lead to a suspicion of ADHD.

The putamen has been associated with the initiation of movements (e.g., Tricomi et al., 2009). However, it was also found to be activated more by reading words and pseudowords than by naming pictures and colours (Oberhuber et al., 2013), the effect being stronger in the left than right putamen, suggesting that it belongs to the reading network. The left putamen was found to predominantly coactivate with left-hemispheric regions that have a direct association with language processes (see Vinas-Guasch and Wu, 2017, for a meta-analysis). Our finding of reduced GM in the left putamen in participants with DD is therefore highly compatible with these results.

The claustrum is a narrow structure located between insula and putamen, the functions of which are not well-known. It is thought to be the most densely connected brain structure, involved in integrating a range of cortical inputs and in segregating attention (Goll et al., 2015). Perhaps this diminished GM volume in claustrum in our dyslexic sample might be related to the observed problems in integrating sensory information (e.g., Widmann et al., 2012) and in regulating attention in DD (see Krause, 2015, for a review).Yet, at this stage these suggestions are highly speculative and need confirmation from further studies.

In the current study, group differences in WM volume were limited to right MTG, hippocampus and precuneus, with dyslexic participants having a smaller volume than controls. MTG (bilaterally) was suggested to be involved in language comprehension (Binder, 2017), and overactivation in right MTG in dyslexic individuals during reading might reflect compensatory functions (meta-analysis of Richlan et al., 2009). Precuneus, in turn, is involved in a wide range of vastly integrated tasks, including episodic memory retrieval and visuo-spatial imagery (Cavanna and Trimble, 2006). Hippocampus, as already discussed, has a central role in memory functions (Buzuki and Moser, 2013). Also some previous studies have shown reduced right-hemispheric WM volumes in DD (e.g., in inferior longitudinal fasciculus, Lu et al., 2016, Banfi et al., 2019) as well as absent rightward asymmetry of the inferior fronto-occipital fasciculus in a subpopulation of DD (Banfi et al., 2019). It could be speculated that the diminished right-hemispheric WM volume of our adults with DD might reflect inefficient compensatory mechanisms, since these participants still had a persistent dyslexia in the adult age.

While our study included 23 dyslexic and 22 control participants, which exceed the minimum group sizes [N (>20)] suggested for neuroanatomical studies on DD (Ramus et al., 2018), even larger sample sizes are needed in the future in order to uncover even finer details of neural deficits associated with DD. Additional limitations of this study are related to the formation of neuropsychological composite scores. The single tests included in the scores could reflect different processes within the composite scores. Although we were not able to conduct a factor analysis for the variables of the current study, the composite variables based on previous theoretical and factor-analytic studies proved to have acceptable internal consistency. Another important limitation is that the two groups were assigned based on subtests included in the technical reading composite score, in addition to one more text reading subtest (section 2.1.). Therefore, the grey-matter group differences and grey-matter volume associations with technical reading should be expected to indicate similar results. The correlations can, in addition to a categorical division of groups, reveal the variance of reading skills within the groups, as well as the direction of the association on a linear scale of reading skills.

### 4.3 Conclusions

The current study evaluated the neuroanatomical basis of DD using rigorous neuropsychological testing and multiple VBM analyses refined to curtail the common limitations. We found reduced GM volumes in adults with DD in left STG and IFG, which is compatible with proposed core nodes of reading (Kujala et al., 2007) and findings in meta-analyses on GM volumes in DD (e.g., Richlan et al., 2013; Ramus et al., 2018). Furthermore, GM volume reductions were found in our DD sample in subcortical structures, which have so far scarcely been studied, including left limbic and basal ganglia areas. WM reductions, in turn, included right temporal and hippocampal structures, as well as right precuneus. Importantly, these brain areas were also correlated with reading accuracy and speed in the whole sample, sharing significant overlap between the two findings, suggesting that these results illuminate the core dysfunctional neural reading network in DD. Next, it would be important to determine which of these volumetric changes in DD underlie the reading deficit and which ones have resulted from, for example, reduced exposure to print. Moreover, as multiple genetic and environmental risks may lead to reading deficits (e.g., Peterson and Pennington, 2015, for a review), a delineation of subgroups of distinct reading deficit profiles from a larger sample would increase our understanding of DD. This, in turn, is vital for developing interventions that are applicable to individuals with various reading deficit profiles.

